# Optimization and Parallelization of Sorting by Interfacial Tension (SIFT) for High-Throughput Metabolic Cell Sorting

**DOI:** 10.64898/2026.03.11.710714

**Authors:** Aria Trivedi, Thomas Mathew, Matthew Shulman, Lakshmi Thangam, Pooja Dubey, Charlotte V. Cohen, Kelsey Voss, Paul Abbyad

## Abstract

A systematic optimization of throughput and operational stability in Sorting by Interfacial Tension (SIFT) is presented. Reducing droplet size and enabling a broader distribution of droplet trajectories increased the number of droplets processed per sorting element, resulting in about a four fold improvement in throughput from 30 to 125 droplets per second. Throughput was further enhanced through device parallelization, with devices incorporating two and four independent sorting regions demonstrated. These configurations distributed droplets evenly across sorting elements that exhibited comparable pH sorting thresholds, indicating similar flow conditions and drag forces within each region. Among the designs evaluated, the two-element configuration provided the optimal balance of throughput, robustness, and simplicity, achieving maximum throughputs of about 250 droplets per second. Throughput and pH sorting thresholds were preserved throughout two hours of continuous sorting. The improved platform was applied to examine the relationship between cellular glycolysis and iron homeostasis at the single-cell level for Jurkat cells, revealing a subpopulation of highly glycolytic cells with significantly elevated iron uptake, consistent with prior reports linking iron regulation and T cell metabolism. Collectively, these advances expand the scale, stability, and biological applicability of SIFT, enabling large-scale functional studies while facilitating the capture of rare and metabolically distinct cell populations.

## Introduction

The separation of cell populations is essential in modern cell biology and medicine. The primary workhorse for this task is Fluorescence-Activated Cell Sorting (FACS), which typically sorts cells via a fluorescence probe. Although expensive, it offers high throughput and unmatched versatility. However, FACS ultimately depends on the availability and specificity of suitable probe molecules to identify target populations, which limits its applicability for some separations. Furthermore, it is not readily amenable to sorting live cells directly based on their secretions or metabolic activity. Label-free microfluidic approaches, including acoustophoresis,^1^ dielectrophoresis^2^ inertial microfluidics,^3,4^ and deterministic lateral displacement ^5,6^, enable inexpensive, high-throughput cell sorting. These techniques separate cells by their size, density, deformability and electrical properties.

Our lab has developed a droplet microfluidic technique, <u>S</u>orting by <u>I</u>nter<u>f</u>acial <u>T</u>ension (SIFT), that allows for the passive and label-free sorting of droplets by pH. The technique differentiates itself by its breadth of applications as it can sort enzymes,^7^ cells^8–10^ and amplified DNA.^11^ As acidification of the extracellular space is linked to a cell’s glycolysis via lactate and proton secretion, it is particularly versatile for the sorting of cells based on metabolic activity. It has been used to separate empty vs. cell occupied droplets, live vs. dead cells, cancer vs. non-cancer cells, naive vs. activated T cells, and cells treated with a pharmaceutical vs. non-treated cells.^8–10^

Despite its versatility, the throughput of SIFT has limited its application to certain applications. Many cellular assays are either facilitated by or require large numbers of cells. Isolating sufficient material for downstream analyses—such as bulk RNA sequencing and mass spectrometry–based metabolomics—typically requires pooling large numbers of cells.^12,13^ Moreover, the ability to sort large numbers of cells is essential for isolating rare populations such as antigen-specific activated effector T cells or circulating tumor cells,which occur at very low abundance yet often exhibit distinctive glycolytic metabolic states.^4,15^

The throughput of our first iterations of SIFT devices was approximately 30 droplets per second.^8,9^ A similar device used to sort amplified DNA droplets also achieved throughputs of tens of droplets per second.^11^ A higher throughput (∼70 droplets per second) was reported in our most recent paper,^16^ which incorporated one of the changes described here, the smaller droplet size. Although other interfacial-tension–based sorting devices have demonstrated higher sorting rates (up to 250 Hz),^17^ direct comparison is difficult because those systems did not sort based on pH or cell glycolysis. The higher throughputs reported in those studies may therefore not reflect differences in the device, but instead could arise from the larger interfacial tension between the specific aqueous samples and the continuous oil phase used, potentially enabling higher flow rates without compromising sorting accuracy.^17^

Here, we describe a systematic increase in the total number of cells sorted by (i) optimizing the droplet size and path, (ii) parallelizing the sorting regions, and (iii) increasing the device run time. Taken together, these modifications increase the sorting throughput by several fold, reaching ∼250 droplets per second, while extending the operational runtime by a factor of six.

As a biological demonstration of the improved device, single-cell intracellular iron levels were measured in activated T cells separated according to high and low glycolytic activity. T cells play a central role in the adaptive immune response, and iron has been shown to enhance glucose metabolism and glycolysis in immune cells, including T helper cells, through activation of metabolic reprogramming pathways such as AKT–mTOR and upregulation of the glycolytic regulator PFKFB4.^18,19^ However, the interplay between glycolysis and intracellular iron during early T cell activation remains poorly understood. We demonstrate here that some highly glycolytic T cells isolated by our device exhibited elevated intracellular iron levels. Improvements in device throughput and robustness significantly broaden the scope of investigations that can be addressed using SIFT.

## Materials and Methods

### Cells

K562 human chronic myelogenous leukemia cells were grown in ATCC-formulated Iscove’s Modified Dulbecco’s Medium (IMDM) and Jurkat Clone E6-1 TIB-152™ Human Acute T-cell leukemia cells were grown in ATCC-formulated RPMI-1640 Medium. Both cell lines were purchased from ATCC. Cells were grown at 37°C in a 5% CO_2_ atmosphere supplemented with 10% fetal bovine serum (HyClone, GE Healthcare Life Sciences, Logan, UT) and 2% v/v penicillin-streptomycin (10 000 units/mL-10 000 μg/ mL) solution (Gibco, Life Technologies Corporation, Grand Island, NY).

### Cell Preparation for on-chip experiments

Jurkat cells were activated with soluble activation complexes (ImmunoCult, StemCell Technologies) following manufacturer protocol and incubated for 24 hours. This initial activation step was not performed for K562 cells. On the day of the experiment, cells were centrifuged, washed and resuspended in buffer, HBBS for Jurkat cells and PBS for K562 cells. To better identify droplet occupancy, K562 cells were labeled by incubating for 30 min at 37 °C and 4% CO2 atmosphere with Calcein AM (Thermo Fischer, Waltham, MA, USA), a viability fluorescent dye. Alternatively for the iron assay on Jurkat cells, cells were labeled with BioTracker Far-red Labile Fe^2+^ Live Cell Dye (MilliporeSigma, Burlington, MA) at 5 µM solution in HBSS for 30 min at 37℃ at 5% CO_2_. Calcein AM was not used when staining with Biotracker to avoid spectral overlap in the fluorescence signals. Subsequently, the cells were washed again and resuspended in on-chip solutions at a cell concentration of 1.0-1.5 × 10^6^ cells/mL, which was determined using a Cellometer Auto T4 Bright Field Cell Counter (Nexcelcom Bioscience LLC, Lawrence, MA). The cell solution is kept dilute to avoid multiple cell occupancy in a droplet. The cell solution was then filtered through a Flowmi 40 µm cell stainter (MilliporeSigma, Burlington, MA) to ensure debris was removed from the solution. On-chip solutions were a 1:1 mix of media and 1.5 mM PBS buffer. The media was prepared without fetal bovine serum (deproteinated media), both solutions were supplemented with 1% w/w Pluronic F-68 (Affymetrix Inc., Maumee, OH), 15% v/v Optiprep solution (Fresenius Kabi Norge AS for Axis-Shield PoCAS, Oslo, Norway) and 0.1 mg/mL pyranine (AAT Bioquest Inc., Sunnyvale, CA). Solution pH and osmolality (determined with Vapro Vapor Pressure Osmometer 5520, Wescor, ELITech Biomedical Systems, Logan, UT) of on-chip solutions were adjusted to physiological values (pH 7.4-7.6; 280−320 mOsmol) prior to experiment. Pluronic F-68 was used to promote droplet stability and cell viability, whereas Optiprep modulated solution density to limit cell sedimentation within the syringe, tubing and droplets.

### Microfluidic Device

Chips with channel depth modulations were fabricated from polydimethylsiloxane (PDMS), utilizing the dry-film photoresist soft lithography technique previously reported by Stephan et al.^20^ This technique facilitates easy prototyping with multilevel designs. The PDMS chip was irreversibly bonded to a glass slide via plasma treatment. To render the internal surfaces of the channel hydrophobic, the channels were treated with Novec 1720 electronic grade Coating (3M, Maplewood, MN) for 30 min at 150 °C. Channel geometry and dimensions for the device with two sorting regions are shown in Figures S1 and S2, respectively, while those for the device with four sorting regions are provided in Figures S3 and S4.

### Droplet Sorting and Measurements

The general use of the sorting device was similar to what has been described previously^8,10^ and is described here for the device with two sorting regions. Briefly, the chip consists of a droplet generator where cells are encapsulated into droplets; an incubation channel enabling a change in droplet pH due to the cells’ metabolism; and a sorting region. Cellular solution was injected into the chip through an aqueous inlet. Via a flow focuser, droplets were generated in 0.1% w/w Picosurf-1 surfactant oil (Sphere Fluidics Limited, Cambridge, United Kingdom) in Novec 7500. An additional oil outlet after droplet generation was set to flow in the opposite direction of the main flow to reduce the amount of oil before the droplets entered the incubator region. This enabled tight packing of droplets within the incubator to ensure the same incubation times for all droplets.^21^ The length of the incubation channel was 10 cm. Incubation time ranged from 3 to 6 minutes depending on the experiment before the channel narrowed and droplets entered the sorting region. At the end of the incubator, oil solution, QX100 droplet generation oil for probes (Biorad, Hercules, CA), entered the chip through two inlets, the QX100 inlet and the Oil Entrainment Inlet (Figure S1). This oil/surfactant combination is called here QX100 for simplicity and consistency with prior publications. Droplets entered the two sorting regions, which both had the same dimensions and flow. The rails, oriented at 45 degrees to the flow direction, allowed sorting droplets by interfacial tension and hence pH. Selected and Unselected populations from both sorting areas are combined before entering the collection outlets.

Flows within the chip were controlled via a computer-controlled syringe pump system (Nemesys, Cetoni, Korbussen, Germany). A very small stir bar (2×2 mm) was placed inside the 0.5mL cell solution syringe. A large external magnet was used to gently mix the solution every 10 minutes to prevent cell sedimentation. PEEK tubing (inner diameter: 180 µm) connected the cell solution syringe to the chip (VICI Precision Sampling, Baton Rouge, LA). Typical flow conditions can be found in the Supplemental Table S1. The temperature of the chip during experiments was maintained at 37 °C using a heating stage with a control module and temperature feedback (CHS-1 heating plate, TC-324C temperature controller, Warner Instruments, Hamden, CT).

On-chip images and videos were taken on an inverted fluorescence microscope (Olympus IX-51) equipped with a 4X objective, a shuttered LED fluorescence excitation source (Spectra-X light engine, Lumencor, Beaverton, OR) and a high-speed camera (VEO-410, Vision Research, Wayne, NJ). The microscope filter cube contained a dual-edge dichroic mirror (Di03-R488/561-t1-25 × 36, Semrock, IDEX Health & Science LLC Rochester, NY) and dual-band emission filter (FF01−523/610-25, Semrock) that enabled transmission of pyranine and Calcein AM fluorescence. The excitation source with individually addressable LEDs was coupled to an Arduino (Arduino LLC, Scarmagno, Italy) to enable rapid alternation between LED colors through simple TTL triggering for determining droplet pH values. Droplets were excited with alternating violet (395 nm BP 25 nm), blue (440 nm BP 20 nm) and green excitation (561 nm BP 14 nm) at a rate of 100 frames per second (33 fps for each color) for pyranine pH measurements. For long sorting experiments, 2 minute videos were taken about every 10 minutes and data was combined.

### Cell Collection

Cell collection and throughput was performed using a workflow first described in Shulman et al.^16^ and summarized in Figure S5. Cells were first sorted as described above and collected into 1mL pipette tips inserted directly into the chip outlets. The pipette tips were prefilled with 300 µL of Novec 7500 to dilute the surfactant found in QX100. Minimizing exposure to this acidic surfactant was found to improve cell viability. 300 µl of HBSS droplets (diameters of 50-200 μm) made in 0.1% w/w Picosurf-1 surfactant oil was also added to the pipette tips. The empty droplets improved cell recovery by ensuring that sorted cell droplets did not collect on the pipette walls and facilitated droplet coalescence. To avoid overflowing the pipette tip over the course of the experiment, oil was removed every 10 minutes from the pipette tip using a long blunt needle syringe. Care was taken during the removal of oil to avoid disturbing the chip or provoking the coalescence of the droplets layered above the oil. After each oil removal, 100 µL 0.1% w/w Picosurf-1 surfactant oil was added to the pipette tip to improve post-sorting cell viability. The pipette tips were capped between transfers to avoid possible contamination.

At the end of the sorting experiment, pipette tips were removed and the oil was drained from the bottom of the tip. Droplets were then collected into microcentrifuge tubes. Droplets were coalesced using a static gun (MILTY Pro Zerostat 3).^22^ To assist in droplet coalescence, Novec 7500 was added to the microcentrifuge tubes during the coalescence process. Once the droplets were coalesced, 20 µL of each cell solution was transferred into individual wells of a 96 well fluorescence plate. To sediment the cells to the bottom of the plate for imaging, the plate was centrifuged at 1500 rpm (g-force 525) for 5 minutes using a TX-100 swinging bucket rotor (Thermo Scientific) at 20 ℃ in a Sorvall Legend XTR Centrifuge.

### Data Analysis

ImageJ software was used for image analysis.^23^ pH values of individual droplets were determined at the end of the incubation channel before droplets entered the sorting region via the ratio of fluorescence intensity from background-subtracted ratio of two color excitation. For pyranine, a calibration curve from fluorescence ratios of blue and violet excitation for droplets of known pH was used to determine pH using a procedure described previously.^9^ Green excitation was used to identify cells labeled with Calcein AM. In one experiment presented here, an elevated rail position within one of the sorting regions led to some droplets missing the rail entirely. In this case, droplets that did not encounter the rail represented less than 5% of the total and were excluded from the analysis. Logistic regression was used to statistically estimate optimal pH thresholds to separate selected from non-selected cells. The pH threshold was defined at a 50% predicted probability of selecting the cell. The standard error of the prediction was used to obtain a 95% confidence interval around that threshold.

Collected cells were imaged with an inverted fluorescence microscope (Olympus IX-50) equipped with a 20X objective, a shuttered LED fluorescence excitation source (Sola SE-II), Lumencor, Beaverton, OR) and a CMOS camera (Orca Flash 2.8, Hamamatsu). Fluorescence images were obtained with long exposure (1.9 seconds) using a red fluorescence cube (Ex: 631 nm BP 28 nm, Dichroic mirror 652 nm, Em: 680 nm BP 42 nm). Debris and irregularly shaped cells were excluded from the analysis. Maximum cellular fluorescence was measured in ImageJ as the dye is expected to be localized in the endoplasmic reticulum based on the manufacturer’s documentation.

## Results and Discussion

SIFT allows for the passive and label-free sorting of cells based on cell glycolysis. The technique leverages a dependence on the droplet interfacial tension with pH in the presence of a surfactant. Cell glycolysis is coupled with a secretion of protons, leading to a decrease in droplet pH. This enables the selection of droplets by interfacial tension and hence single-cell glycolysis.

Figure 1a shows the basic chip geometry of the current SIFT device, which integrates cell encapsulation, incubation, and sorting into a single continuous-flow inline platform. In many other droplet sorting devices,^24,25,17^ these steps are decoupled: cells are encapsulated into droplets in one device, incubated off-chip and then droplets are injected into another device for sorting. Integrating these steps within a single device eliminates user intervention between stages and reduces potential failure points associated with droplet collection and reinjection. It also simplifies setup, as only one chip needs to be prepared. Briefly, the device operates as follows. Cells are encapsulated into droplets. Cell concentration is kept low to avoid multiple cell occupancy in a droplet. Droplets then flow through a long serpentine incubation channel. During this time, cell glycolysis will lead to an acidification of the droplet. Lastly, droplets enter the cell sorting step. Figure 1b shows a detailed view of the sorting geometry (channel geometry and dimensions of sorting region are provided in Figures S1 and S2, respectively). In this diagram, a channel geometry with two parallel sorting regions is presented.

**Figure 1.**
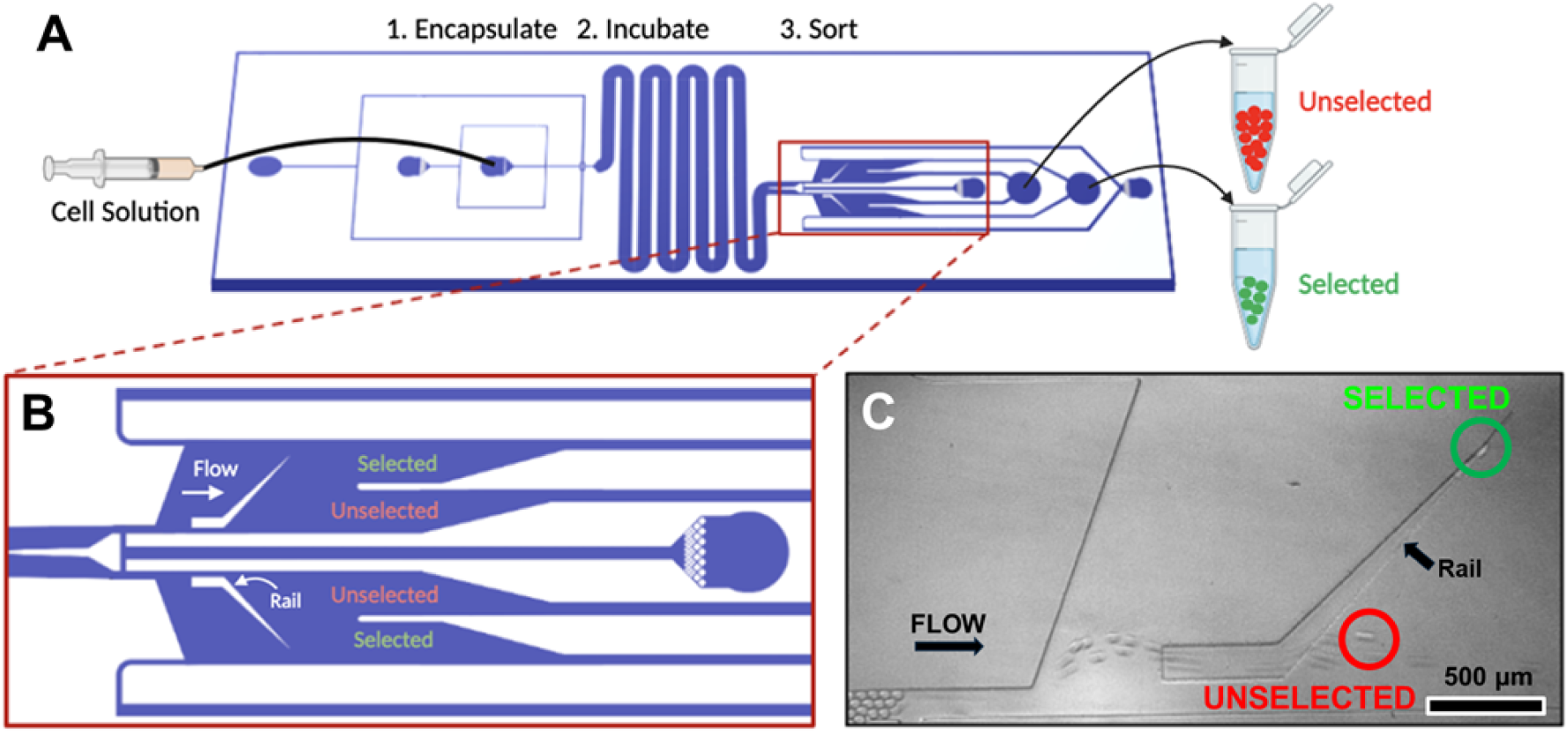
a) Schematic of SIFT chip device for sorting a population of cells with low (red) and high (green) glycolysis. Created in BioRender.com. b) Chip geometry of device with two sorting regions c) Image of sorting showing a droplet containing a cell with high glycolysis ride the rail laterally up (circled in green). Empty droplets or those containing cells with low glycolysis (circled in red) are only slightly deflected by the rail.

As droplets enter the sorting step, a carrier oil solution (QX100) is introduced, which exhibits an inverse relationship between droplet pH and interfacial tension. Droplets, flattened by the top and bottom of the channel, enter the rail region of increased channel height. The droplets are less confined in the rail and therefore adopt a more spherical shape to minimize their surface energy. Droplets of high pH and hence low interfacial tension are pushed off the rail by the oil entrainment flow and directed to an Unselected chip outlet (Figure 1c). Droplets of low pH, and hence high interfacial tension, remain in the rail as the entrainment flow is insufficient to reconfine the droplets. The droplets follow the rail, oriented 45 degrees relative to the direction of flow and exit the tapered rail near the top. These droplets are directed towards the Selected chip outlet. In this way, cells with high or low levels of glycolysis are separated and directed towards different chip exits for recovery.

Early iterations of the SIFT device^8,9^ had throughput of approximately 10-30 droplets sorted per second. The device could be run continuously for 10-20 minutes. As most droplets are empty (typically 1 in 20 or 30 droplets contain a cell) this effectively meant the sorting of several hundreds cells. Here we describe a series of optimizations, presented in separate sections covering droplet size and path, parallelization, and stability. These optimizations lead to typical throughputs of 200-250 droplets per second and stability over 1-2 hours. Collectively, these improvements enable the sorting of several thousand cells. The device is also simpler, requiring 5 independent syringe pumps rather than six of previous devices.

### OPTIMIZING DROPLET SIZE AND PATH

A direct way to increase droplet throughput is by increasing the number of droplets that are sorted by an individual sorting element. Several changes were made to increase the sorting throughput. The first was to decrease the droplet size from 70-90 µm to 40 µm. This was achieved by decreasing the nozzle width of the flow focuser from 50 µm to 30 µm.

Using smaller droplets have several implications for both the function and the sorting in the device. The smaller volume means that droplets change pH quicker due to cell glycolysis. Consequently, the incubation time could be short, 3-6 minutes, while still maintaining a pH change suitable for sorting. This allows faster overall flow rates in the incubator. Moreover, due to the smaller size, the overall number of droplets in the incubator is increased. This results in more droplets reaching the sorting step and ultimately sets the upper limit for the sorting rate.

In previous iterations of the device, an effort was made to ensure that droplets entered the sorting region one by one and followed a similar path (Figure 2a). A similar approach is seen in other papers.^11^ This avoided droplet collisions that may impact specificity. To ensure well spaced droplets, an additional outlet (labeled Droplet Outlet in bottom left in Figure 2a) with a negative flow was applied to remove droplets before entering the sorting region.^8,9^ In most cases, approximately half of the droplets were removed from the device prior to sorting. This throttled the number of droplets entering the sorting region ensuring a consistent droplet path. Although this approach yielded strong pH specificity, it ultimately constrained the device’s throughput.

**Figure 2.**
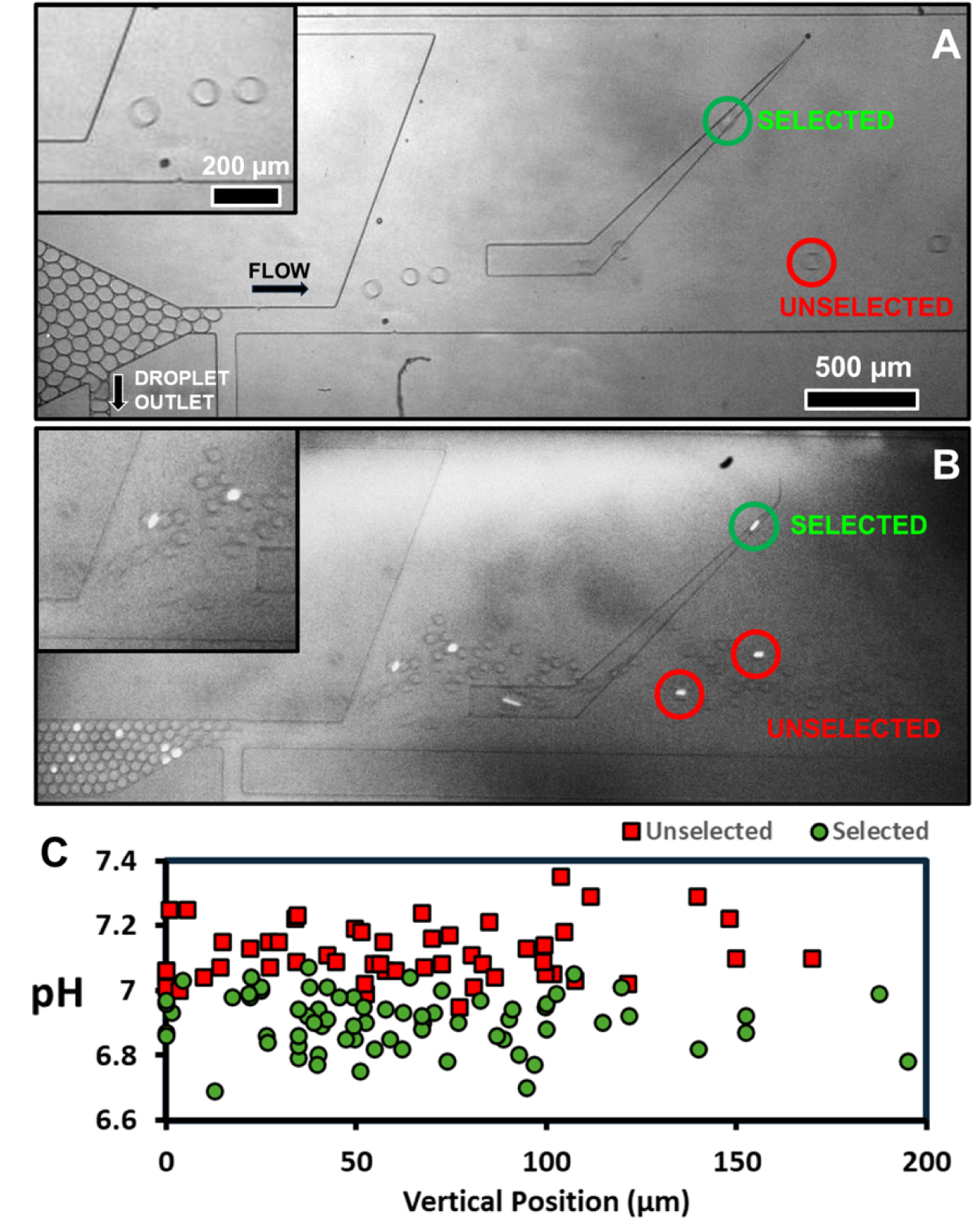
a) Image of sorting region for previous SIFT sorting devices. An Unselected droplet (circled in red) and Selected droplet (circled in green) are highlighted in image. Large droplets (70-90 µm diameter) follow similar path in sorting region. Cells appear as bright spots due to fluorescent labeling. Inset: (upper left) shows an enlarged view of the beginning of the sorting region. b) Image of sorting region of current SIFT sorting device. Smaller droplets (40 µm diameter) follow different paths in sorting region. c) pH of Unselected droplets (red squares) and Selected droplets (green circles) plotted as a function of the maximum vertical position reached by the droplet in the sorting region.

In contrast, using smaller droplets enabled a different strategy: more droplets could be pushed into the sorting region at the end of the incubator. In this case, droplets scatter from the outlet and take different paths in the sorting region (Figure 2b). Based on the droplet dimensions and available paths, roughly three droplets could be spaced to flow laterally in parallel. However, in practice, the droplets were typically staggered. In this new design, all droplets enter the sorting region and no droplets are discarded. This change was found to increase device stability while increasing throughput. It also reduced the total number of syringe pumps, from six to five, necessary to operate the SIFT device.

The pH specificity can be studied as a function of the different paths. Figure 2c shows the selection of droplets based on the maximum lateral, or vertical, position on a droplet’s path. A pH threshold is defined as the average pH of droplet selection and is determined by a fit to a logistic regression. By comparing the boundary between Selected and Unselected droplets, it is observed that the threshold is largely maintained regardless of path. The threshold for all droplets in Figure 2c was determined to be 7.02 ± 0.01 (Figure S6). When the threshold was instead evaluated at different vertical positions (0–50 μm, 50–100 μm, and >100 μm), the corresponding thresholds were 7.03 ± 0.01, 6.99 ± 0.02, and 7.03 ± 0.02, respectively (Figure S7). This indicates that droplet path or interdroplet collisions do not appear to substantially impact pH threshold. As a result of these changes, 100-125 droplets can be sorted by a rail. It is worth noting that the travel time required to reach the rail should be approximately the same for all droplets. Upon exposure to QX100, droplets gradually acidify, leading to a concurrent increase in interfacial tension.^16^ Consequently, a droplet that takes substantially longer to reach the rail may be selected despite having a higher initial pH. Thus, the current device is expected to exhibit slightly lower specificity compared to systems in which droplets follow identical paths. However, many applications can tolerate this modest decrease in specificity in exchange for the several-fold increase in throughput achieved by the present design.

### INCREASING THROUGHPUT VIA PARRALLIZATION

SIFT is a passive technique that requires no active sorting components or detectors. As a result, it is particularly well-suited for parallelization of sorting modules, since it does not increase overall cost or system components. Previous versions of SIFT devices had a single sorting region and we present here devices with either 2 or 4 sorting regions.

Figure 3a and Video S1 demonstrates a chip geometry with 2 sorting regions that are mirror images of each other. The droplets exiting the incubator are distributed evenly between the two sorting regions (Figure 1b). Immediately prior to entry in the sorting regions, QX100 was introduced in the device positioned between the two sorting regions (Figure 1b). A QX100 inlet, dubbed Oil Entrainment Inlet, was positioned in the lateral extremities of the sorting regions. This outlet allowed a control of the oil flow in the sorting region and thus provided a user defined parameter to adjust the pH threshold. The sorting rail deflects the lateral path of droplets containing cells with high glycolysis. Sorted droplets from the two regions are merged into common Selected and Unselected outlets, preserving the same number of outlets as in earlier versions of the device.

**Figure 3.**
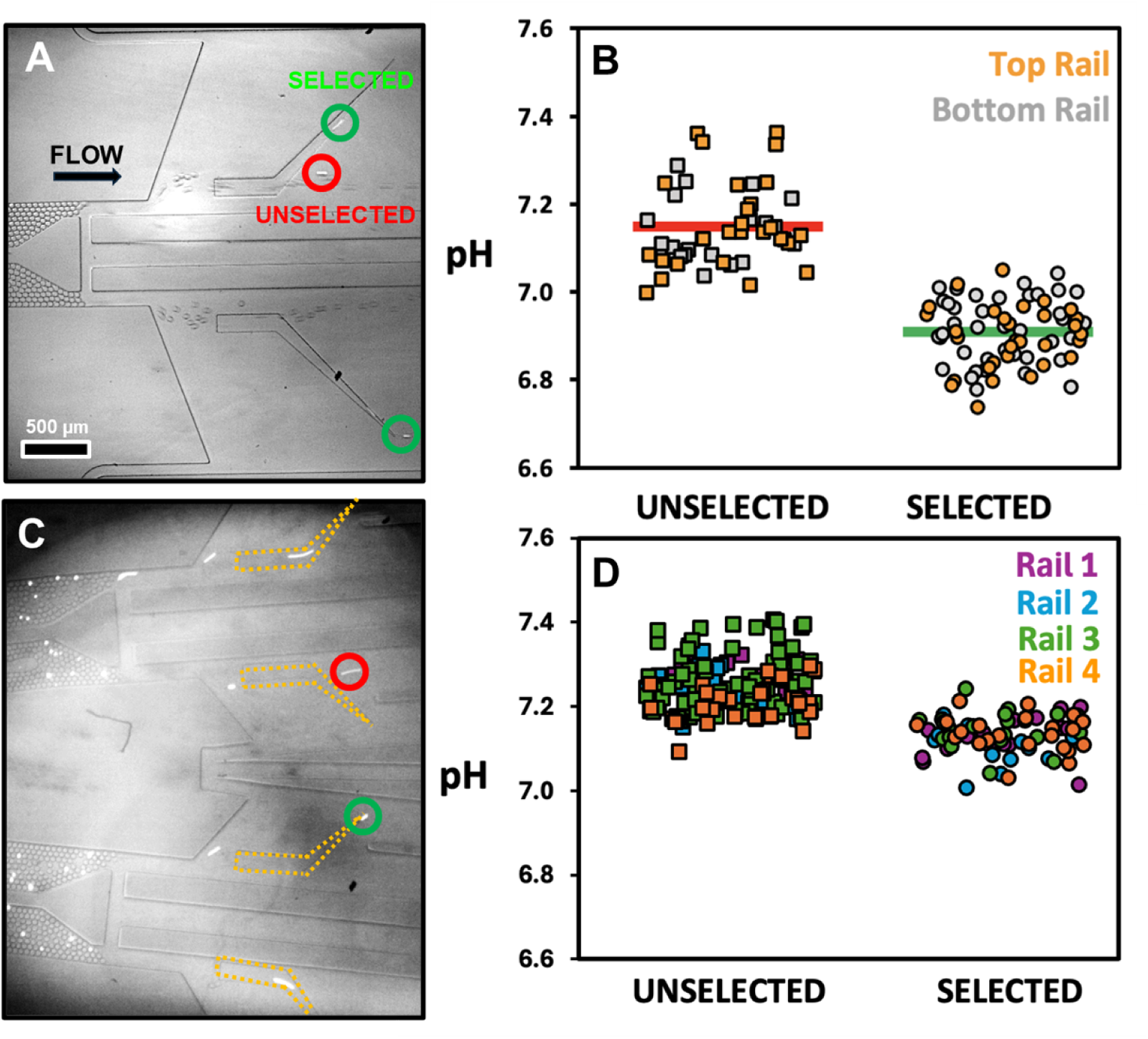
Image of sorting region for a device with two sorters. An Unselected droplet (circled in red) and Selected droplets (circled in green) are highlighted in image. K562 cells appear as bright spots due to fluorescent labeling. b) pH of droplets containing cells for Selected and Unselected populations for the top and bottom rail. Average pH value indicated by horizontal bar. c) Image of sorting region for a device with four sorters. Rails are outlined with dashed yellow lines to enhance visibility in the image. d) pH of droplets containing cells for Selected and Unselected populations for the four rails. Rails are numbered from top to bottom.

The throughput of the device was determined to be 100-125 droplets per second for each of the sorting regions or approximately 200-250 droplets per second in total. This represents an increase of about eight fold from previous chip designs. Figure 3b shows the droplet pH of the two sorting regions. The top and bottom rails exhibit similar thresholds of 7.02 ± 0.01 and 7.04 ± 0.01 (95% confidence interval), respectively (Figure S8). For the bottom rail, the two populations show no overlap, preventing direct determination of the error from the fit; instead, the uncertainty is estimated from the closest data points between the two populations. This similarity in threshold indicates that doubling the number of sorting regions can be obtained without compromising pH specificity. The threshold however is heavily dependent on the rail position^16^ and thus care must be taken to ensure that the rails are positioned in identical positions within the sorting region. The lateral position of the rail is also important to ensure that all droplets encounter the rail. If the rail is positioned too high, some droplets may miss the rail entirely which would impact the specificity of the Unselected population of cells.

Figure 3c and Video S2 shows a device with four sorting regions. To avoid expanded significantly the chip width, the sorting region was reduced by half from a lateral width of 1500 µm to 750 µm. Figure 3d shows similar pH thresholds for all four sorting rails, arranged from top to bottom, with pH values of 7.20 ± 0.01, 7.16 ± 0.01, 7.17 ± 0.01, and 7.16 ± 0.01 for rails 1–4, respectively (Figure S9). However, the overall sorting throughput of the device with four sorting regions (250 droplets per second) was comparable to that of the device with two sorting regions, likely because the reduced channel width constrained the number of available droplet paths. Additionally, a single-layer device cannot allow the merging of outlets belonging to the same population, whether selected or unselected, into a single outlet. This can only be achieved either off-chip or in a two-layer device,^26^ but doing so increases device complexity. Although this design demonstrates the feasibility of further multiplying the sorting regions, the associated complexity and throughput did not justify its use. Consequently, the two-sorting-region device was deemed a more practical device and used for all applications and discussion below.

### INCREASING CELL COLLECTION THROUGH EXTENDED RUN TIMES

The operational stability of microfluidic devices over extended runtimes—though often unreported—is a critical performance metric for practical applications. As an inline device, the number of sorted cells is proportional to the overall run time. However, this requires a stable device over time with a steady input of cells. A particular challenge associated with long run times is the tendency of cells to sediment in the syringe and tubing before reaching the microfluidic device. This sedimentation is exacerbated by the low flow rates of the cell suspension in our device (typically 0.3 μL/min, Table S1). A density modulator (OptiPrep) is used to match cell and solution densities. While this reduces cell sedimentation, the number of cells entering the device decreases substantially after approximately 20 minutes. Optimizing the concentration of Optiprep can help alleviate but does not eliminate the problem.

To ensure a constant flow of cells over the entire device run time, a syringe was used with a small magnetic stir bar placed inside. About every ten minutes, the magnetic stir bar inside the syringe was moved back and forth using a larger external magnet to resuspend the cells. Furthermore, PEEK tubing with a small inner diameter (0.18mm) was used to connect the syringe to the device. The small inner diameter limited cell sedimentation during the approximately 25-minute residence time in the tubing. Together, these modifications maintained a consistent influx of cells into the device, allowing for experiments of extended duration.

Figure 4 summarizes the temporal stability of the device, based on measurements obtained at discrete time points over several hours. As shown in Figure 4a, the droplet sorting rate remains stable at approximately 200–225 droplets per second over the entire 4 hour runtime. Figure 4b shows the number of cells sorted per second at different time points, revealing a decrease in cell entry after approximately 2 hours of operation.The decrease in cell entry after two hours may result from gradual cell sedimentation, aggregation, or subtle changes in viability during extended runtimes. Despite this reduction, the pH sorting threshold remains consistent over the 4 hours (Figure 4c and Figure S10), demonstrating that the long-term operational stability of the sorting device remains robust. The long-term stability is critical when working with precious or limited samples and represents a key requirement for translating the technology to a commercial device.

**Figure 4.**
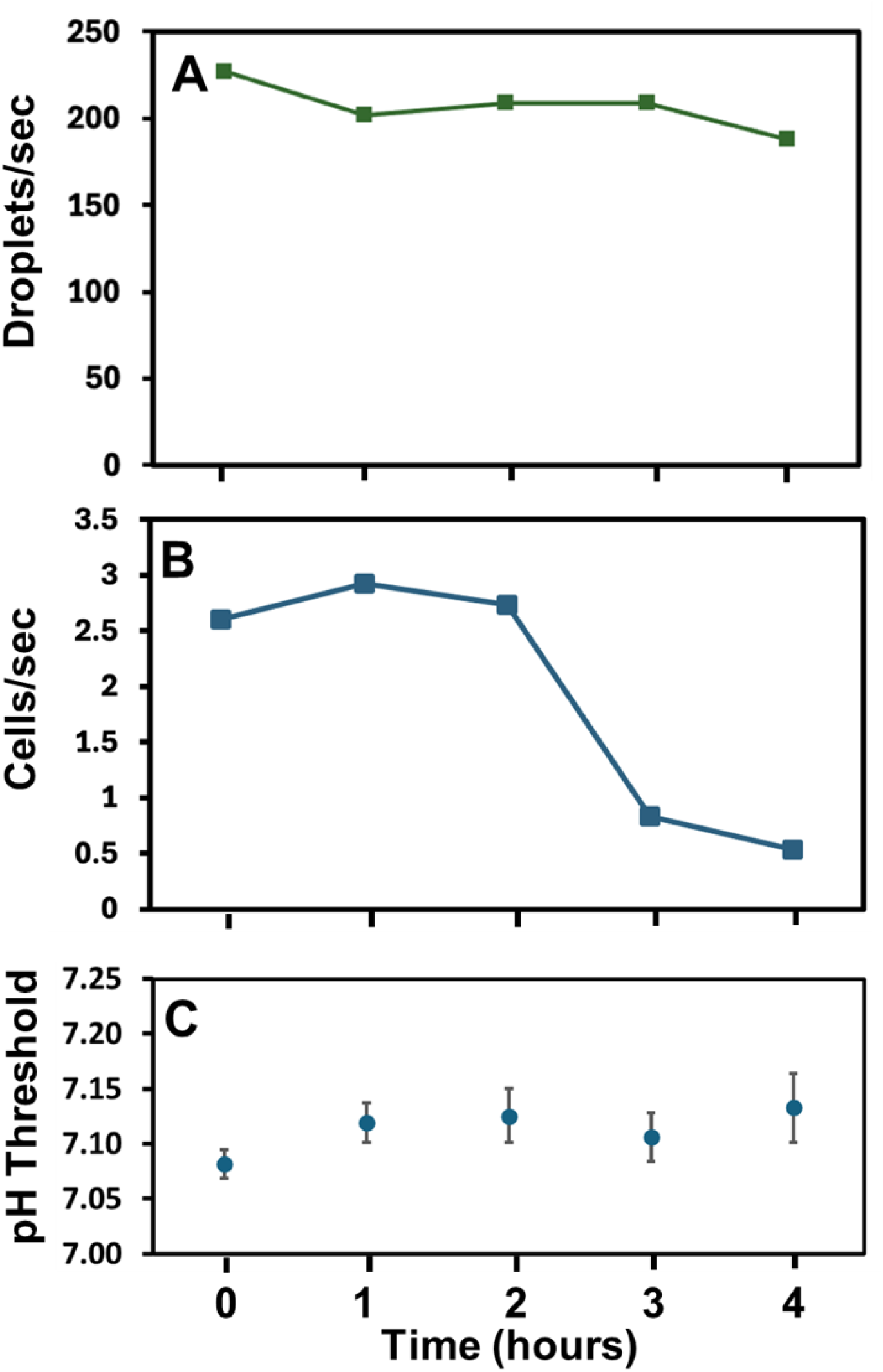
Sorting rate and threshold for a continuous run. All values were determined from 2 minutes of sorting data collected at the respective time points a) Rate of droplet sorting at different time points for the SIFT device. b) Rate of cell sorting. c) pH threshold of sorting. Error bars represent standard error.

### APPLICATION TO CELLULAR IRON UPTAKE

As a demonstration of the sorting capacity of the device, the relationship between glycolysis and iron uptake was explored in activated Jurkat T cells. Iron plays an important role in T cell proliferation and differentiation, in part through its involvement in mitochondrial metabolism.^27^ Iron supports T cell activation by enabling mitochondrial respiration, anabolic metabolism, and redox-dependent signaling.^27,28^ Activated T cells upregulate iron uptake to meet the increased metabolic demands associated with proliferation and effector differentiation.^19^ As activated T cells differentiate to perform specific cellular functions, iron availability and regulation influence functional outcomes. Previous work has shown that iron uptake drives T cells into a proinflammatory phenotype via sped-up metabolic rates,^19^ while hypoferremia, or low iron, counteracts the development of adaptive immunity.^29^ However, the underlying mechanisms by which iron regulation shapes metabolic heterogeneity and functional diversity at the single-cell level remain poorly understood, motivating approaches that can directly link iron uptake to metabolic state in individual cells.

Prior to sorting, activated Jurkat T cells were incubated with a fluorescent iron sensor (BioTracker Far-red Labile Fe2+ Live Cell Dye). Figure 5a shows the sorting of droplets containing activated T cells using the device presented earlier containing two sorting regions with an Selected and Unselected droplet indicated in green and red, respectively. The overall throughput was about 240 droplets per second. Figure 5b shows the pH of individual droplets of the Selected and Unselected droplets. The pH threshold was determined to be 7.29 ± 0.01 (Figure S11). In this case the error is an estimate, as there was no overlap between the two populations. Following sorting, cells were recovered by droplet coalesence using a protocol (Figure S5) described in Shulman et al.^16^ and imaged on a fluorescent microscope. Max cell intensity rather than the cellular average was measured as the labile iron dye is expected to be localized in the endoplasmic reticulum. Figure 5c show the Biotracker single-cell fluorescent intensity of the Selected and Unselected populations. While some Selected highly glycolytic cells share similar iron uptake to low glycolytic cells, there are a significant number of highly glycolytic cells that show increased iron uptake. Selected cells exhibited a 2.27-fold higher maximum cell intensity compared to the low-intensity Unselected population. A Welch’s two-sample t-test confirmed a significant difference between the two populations. A Welch’s test was used because the populations exhibited unequal variances. As single-cell measurements often produce skewed distributions due to rare high-intensity events, the data was log-transformed prior to statistical analysis to better approximate normality. Overall, Figure 5 reveals that glycolysis and iron homeostasis are correlated for a subset of cells. This experiment brings a new approach of examining this relationship at a single cell level allowing for the exploration of cells that are far from the average. This association can be further explored at the single-cell level to elucidate potential mechanisms underlying the correlation between glycolysis and iron uptake.

**Figure 5.**
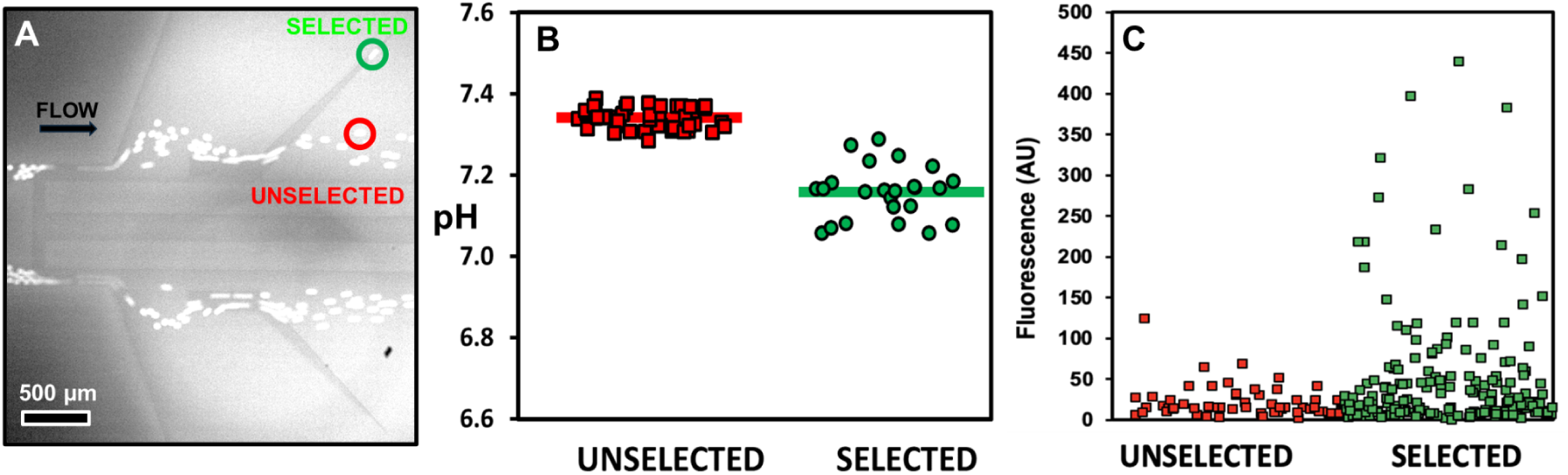
a) Image of sorting region showing Unselected Droplet (circled in red) and Selected droplet (circled in green). b) pH of Selected and Unselected cell populations. Average pH value indicated by horizontal bar. c) Maximum cellular fluorescence intensity of BioTracker Far-red Labile Fe^2+^ Live Cell Dye for the Unselected and Selected cell populations. The Selected and Unselected populations were statistically different (Welch’s t-test, (t(148.1) = −3.13, p = 0.002).

## Conclusions

A systematic improvement of the throughput and stability of the SIFT device is presented. The use of smaller droplets that take a breadth of sorting paths enabled a substantial increase in the number of droplets treated by a single sorting element. This change allowed an increase in throughput of approximately a factor of four from 30 droplets per second to 125 droplets per second. Sorting throughput is further increased by parallelization, and the use of two and four independent sorting regions are demonstrated. These devices distribute the droplets within the sorting elements. The elements are shown to display similar pH sorting thresholds. As drag force is a determinant component in sorting, this indicates similar flow conditions are achieved in the different sorting regions. The device with two sorting elements was found to offer best results with respect to throughput, robustness and simplicity. The sorted populations from the two sorting elements can be easily combined at the outlet. This device had a maximum throughput of about 250 droplets per second. By minimizing cell sedimentation in the syringe and tubing this device was shown to be stable for 2 hours of runtime. Notably, because the system relies on a single device, preparation requires only about 30 minutes, after which it can be operated for extended periods to sort the number of cells needed for the desired application.

Further improvements in throughput are likely possible. Higher throughputs may be possible by using even smaller droplets, however optimal flow and rail position would be needed to maintain reliable sorting. As droplets acidify while flowing downstream from exposure to QX100, their interfacial tension increases. Positioning the rail further downstream than presented here could therefore allow the device to operate at even higher sorting rates. A massively parallel architecture is also conceivable; however, implementing such a design is not trivial due to constraints inherent to the SIFT device. In particular, the QX100 surfactant can only be introduced immediately prior to sorting, which complicates the device layout and limits where parallelization can be implemented.

The improved device was used to study the relationships between cell glycolysis and iron homeostasis, at a single-cell level. The results show that there is a subset of cells that show significant higher iron uptake within the the highly glycolytic cells. These findings support previous research of the interplay between iron homeostasis and T cell metabolism.^28^ This work brings a novel approach to examining iron homeostasis through the lens of single-cell metabolism. The performance improvements significantly expand the investigative capacity of Sorting by Interfacial Tension by supporting biological studies requiring greater scale while enabling the capture of rare and metabolically distinct cell populations.

## Acknowledgments

TM acknowledges generous funding from the Beckman Scholar program. AT acknowledges funding from the Clare Boothe Luce Scholar program. KV was supported by the Breakthrough T1D (formerly the Juvenile Diabetes Research Foundation) Advanced Postdoctoral Fellowship. PA was supported for this project by a National Science Foundation Career Award and the Henry Dreyfus Teacher-Scholar Awards Program.

## Notes

### Competing Interest Statement

The authors have declared no competing interest.

