## Supplemental Info for "Optimization and Parallelization of Sorting by Interfacial Tension (SIFT) for High-Throughput Metabolic Cell Sorting"

### Supplemental Information

**Table of Contents:**

**Video Caption……………………………………………………...………..…..Page S-2**

**Figure S1…...…………………………………………………………...…..Page S-3**

**Figure S2…...…………………………………………………………...…..Page S-4**

**Figure S3…...…………………………………………………………...…..Page S-5**

**Figure S4…...…………………………………………………………...…..Page S-6**

**Figure S5…...…………………………………………………………...…..Page S-7**

**Figure S6…...…………………………………………………………...…..Page S-8**

**Figure S7…...…………………………………………………………...…..Page S-9**

**Figure S8…...…………………………………………………………...…..Page S-10**

**Figure S9…...…………………………………………………………...…..Page S-11**

**Figure S10..…...……………………………………………………...…..Page S-12**

**Figure S11…..…………………………………………………….....…..Page S-13**

**Table S1…......………………………………………………………………..Page S-14**

**Video Caption:**

**Video S1. SIFT device with two sorting regions:**
In this video, two sorting regions are used to increase droplet sorting throughput. K562 cells appear as bright spots due to fluorescent labeling. Droplets with cells with high glycolysis are deflected laterally to the Selected outlet. Empty droplets or those containing cells with low glycolysis are only slightly deflected by the rail and are directed to the Unselected outlet. Video slowed by 3X.

**Video S2. SIFT device with four sorting regions:**
In this video, four sorting regions are used to increase droplet sorting throughput. K562 cells appear as bright spots due to fluorescent labeling. Video slowed by 3X

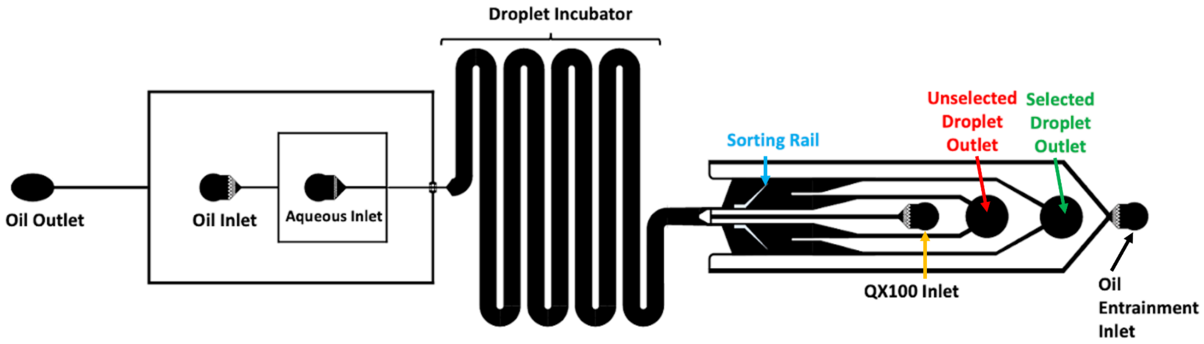


**Supplemental Figure S1. SIFT device with two sorting regions channel geometry.**

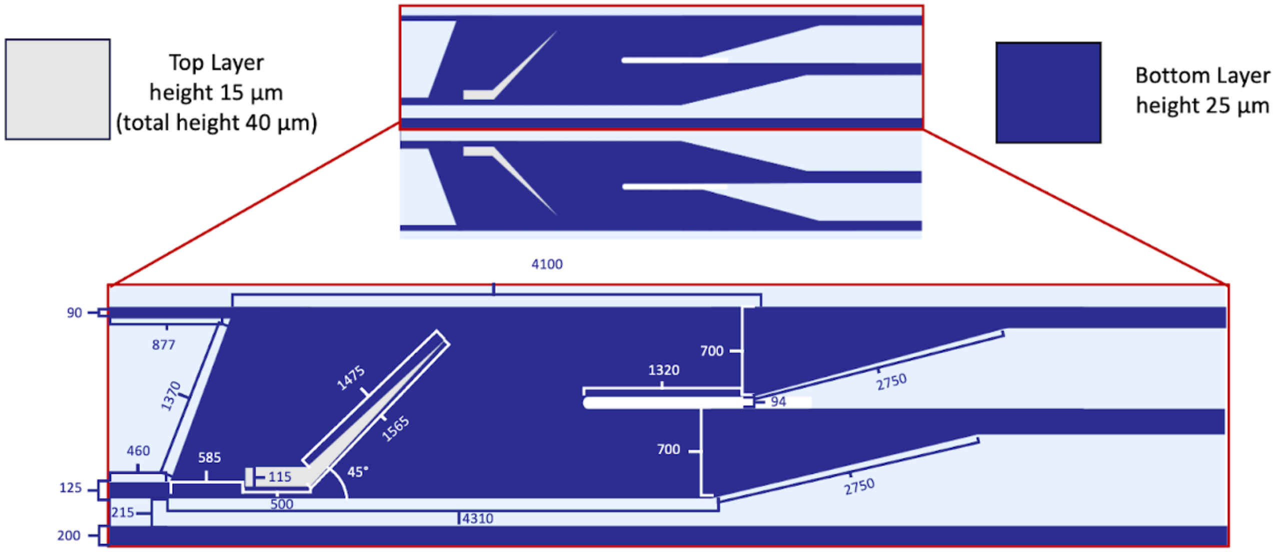


**Supplemental Figure S2. Sorting Area Dimensions for device with two sorting regions.** One set of sorting area measurements are shown as both sorting areas have the same dimensions. The rail position is approximated as the exact position is layered by eye during the microfabrication process.

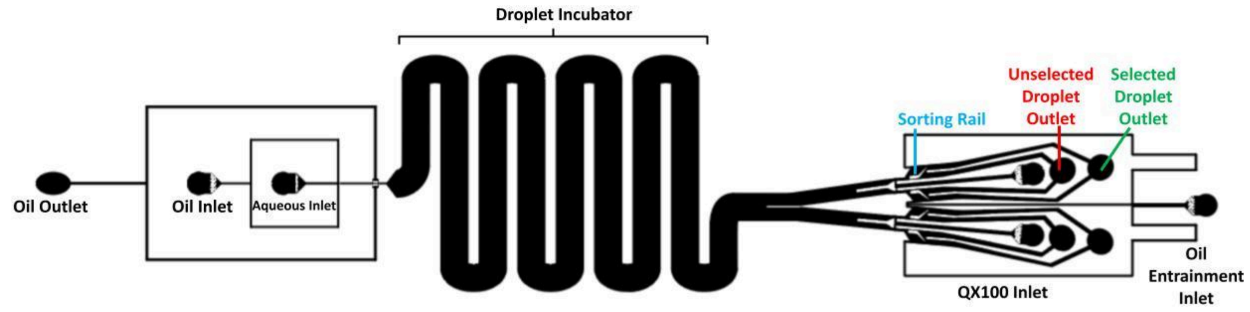


**Supplemental Figure S3. SIFT device with four sorting regions channel geometry.**

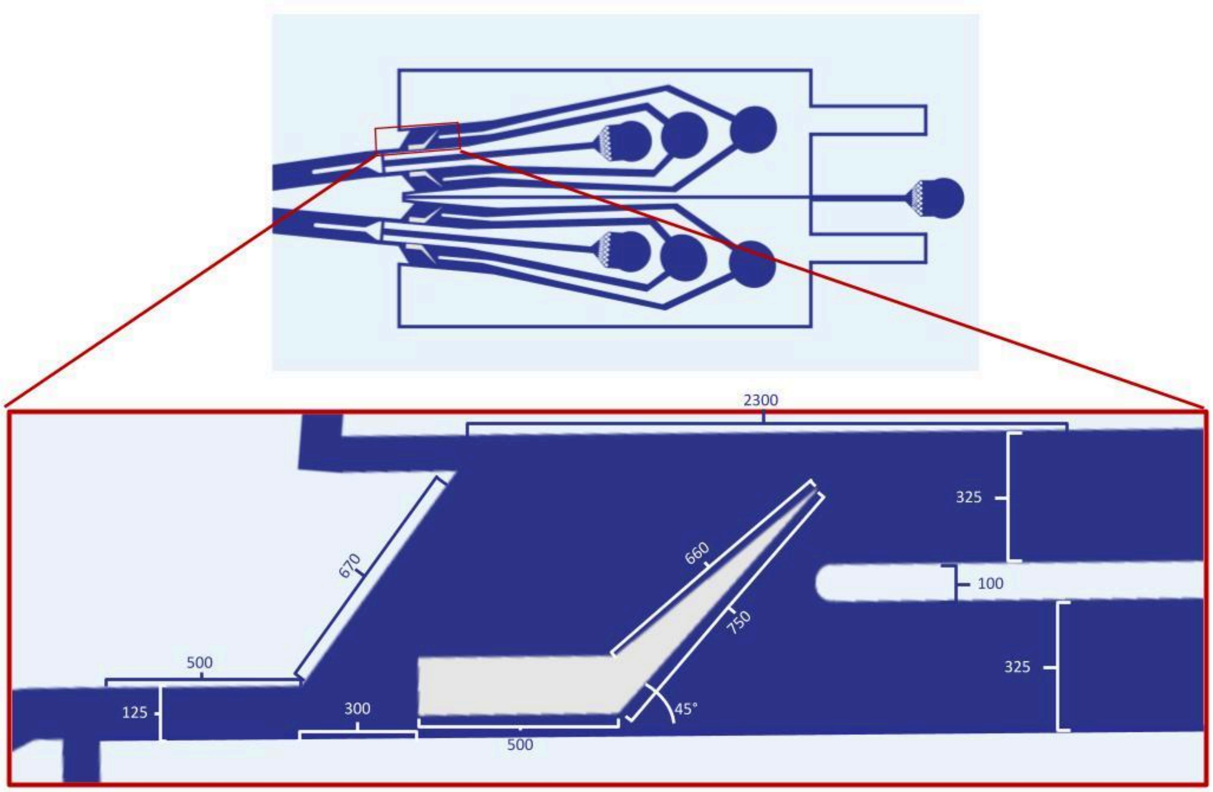


**Supplemental Figure S4. Sorting Area Dimensions for device with four sorting regions.** One set of sorting area measurements are shown as all sorting areas have the same dimensions. The rail position is approximated as the exact position is layered by eye during the microfabrication process.

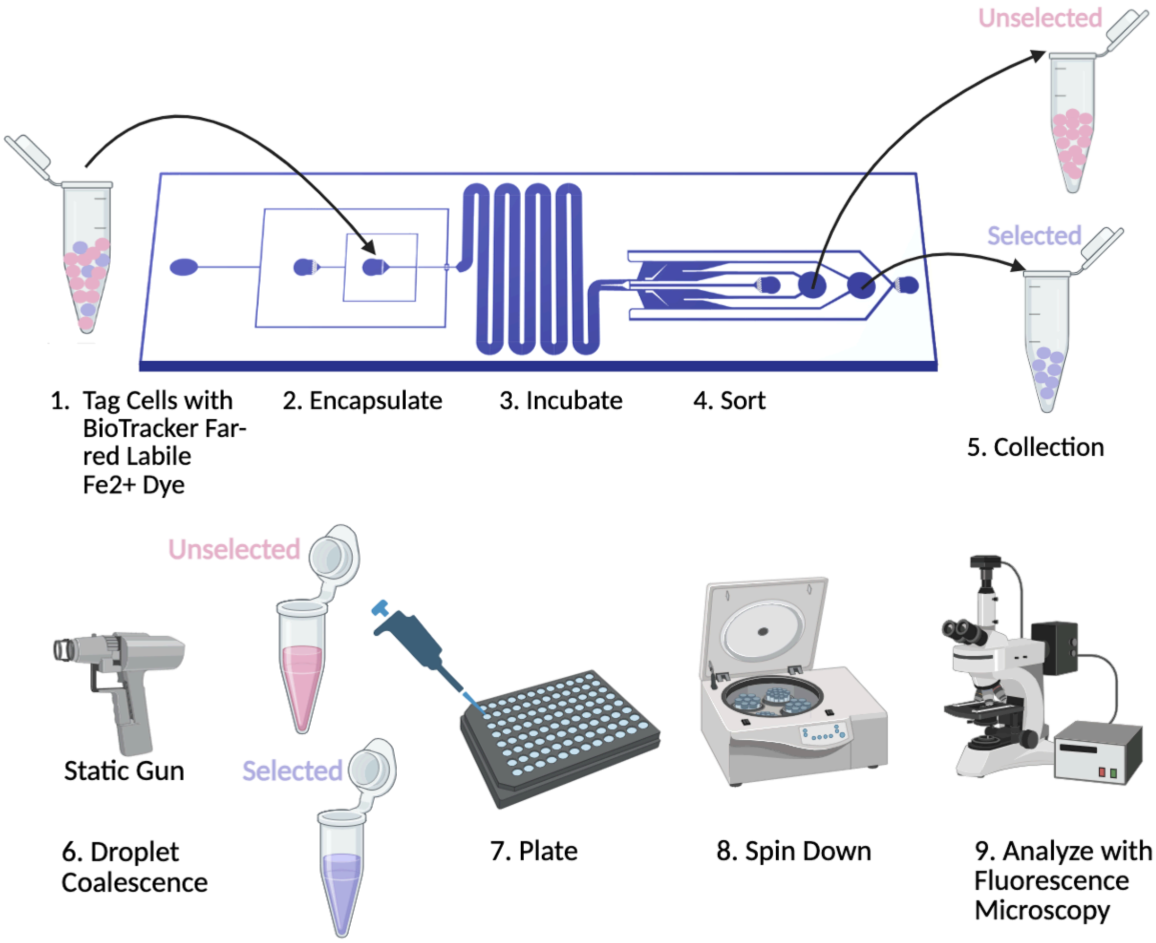


**Supplemental Figure S5.** Workflow for cell collection. Created in BioRender.com.

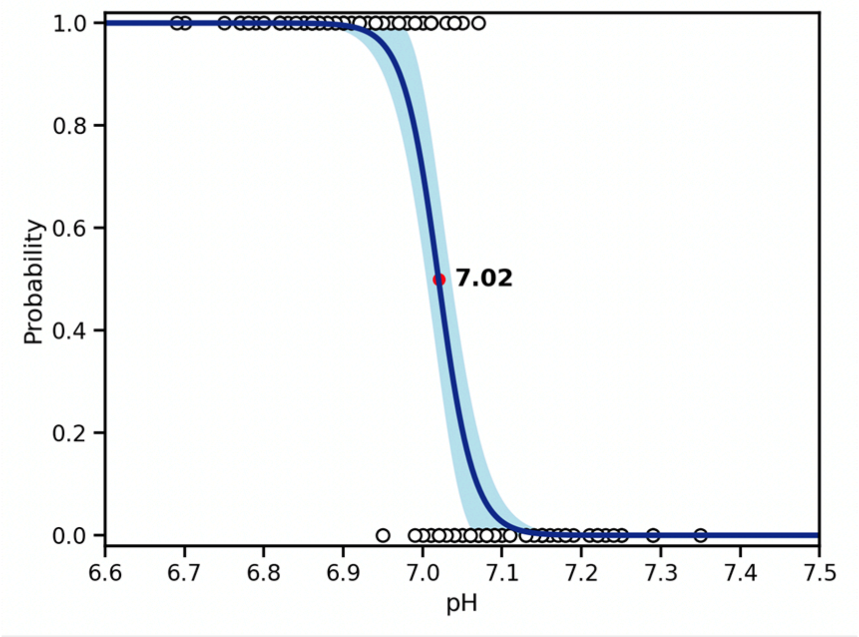


**Supplemental Figure S6.** Logistic regression fit of binary selected/unselected data with pH. pH thresholds are indicated on graph and represent where there is equal probability that droplets are selected or unselected. The 95% confidence limit is indicated in light blue.

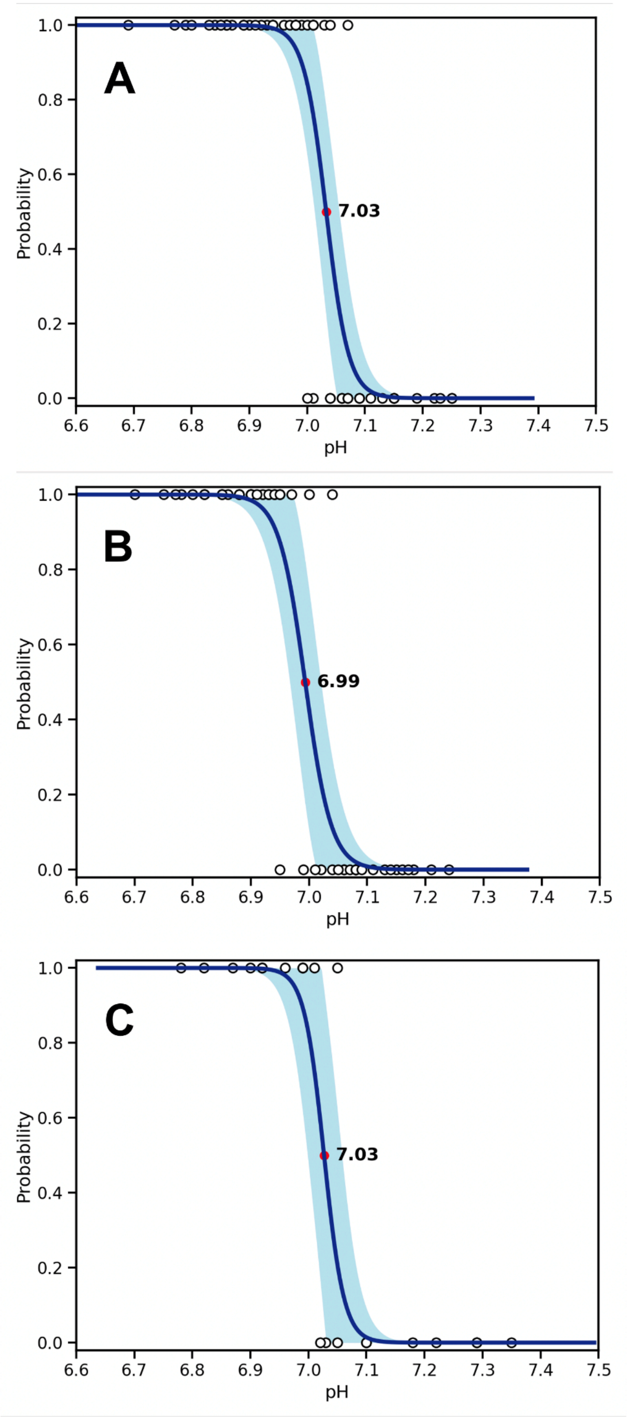


**Supplemental Figure S7.** Logistic regression fit of binary selected/unselected data with pH for a (**A**) maximum lateral position 0–50 μm (**B**) 50–100 μm (**C**) >100 μm.

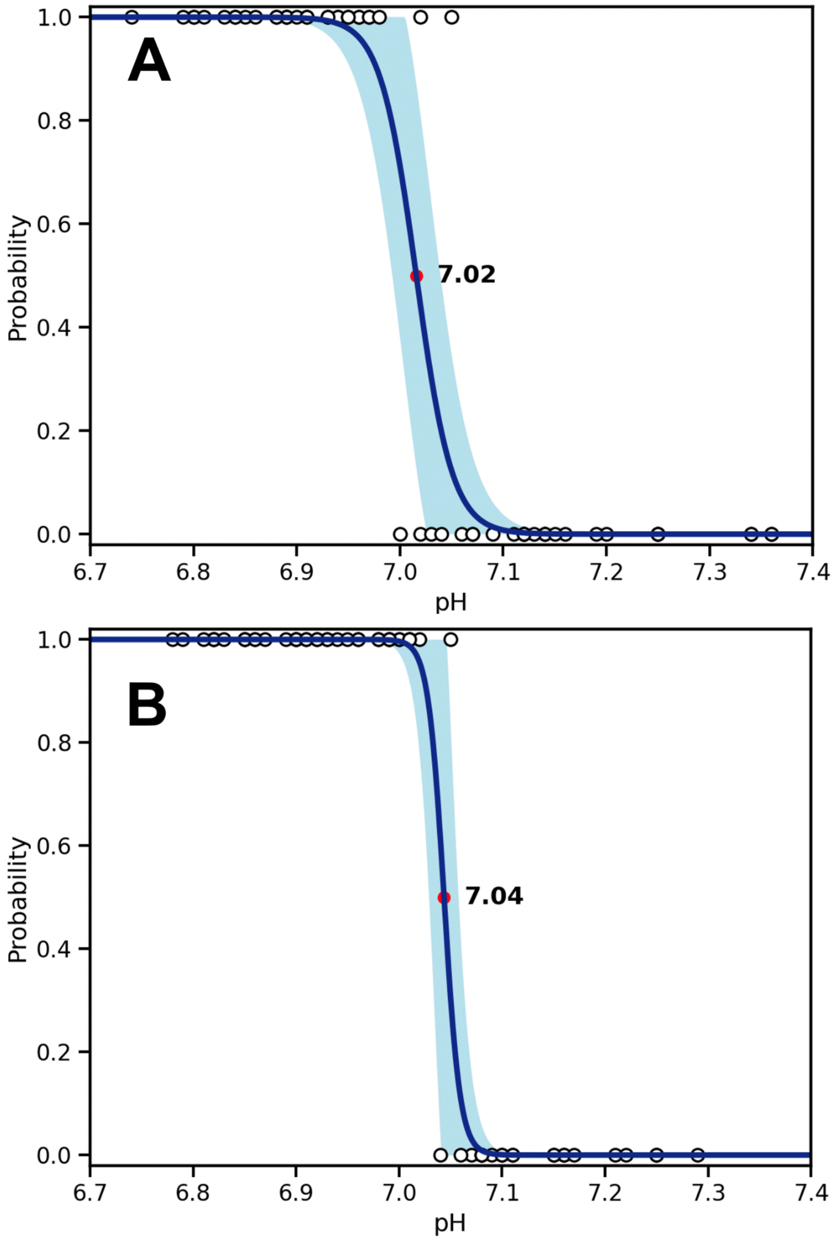


**Supplemental Figure S8.** Logistic regression fit of binary selected/unselected data with pH for a device with two sorting regions. (**A**) Top Rail (**B**) Bottom Rail.

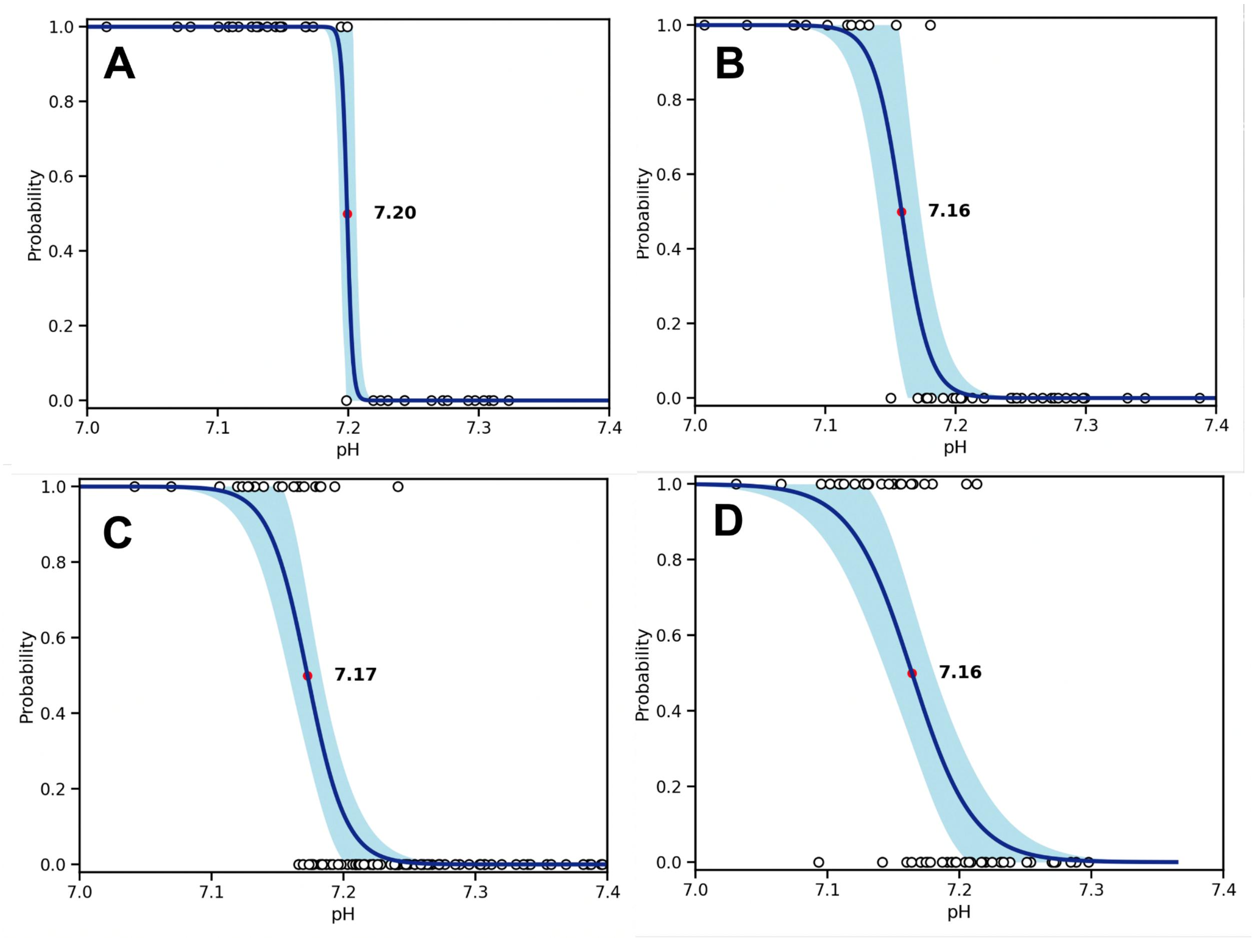


**Supplemental Figure S9.** Logistic regression fit of binary selected/unselected data with pH for a device with four sorting regions. (**A**) Rail 1 (**B**) Rail 2 (**C**) Rail 3 (**D**) Rail 4.

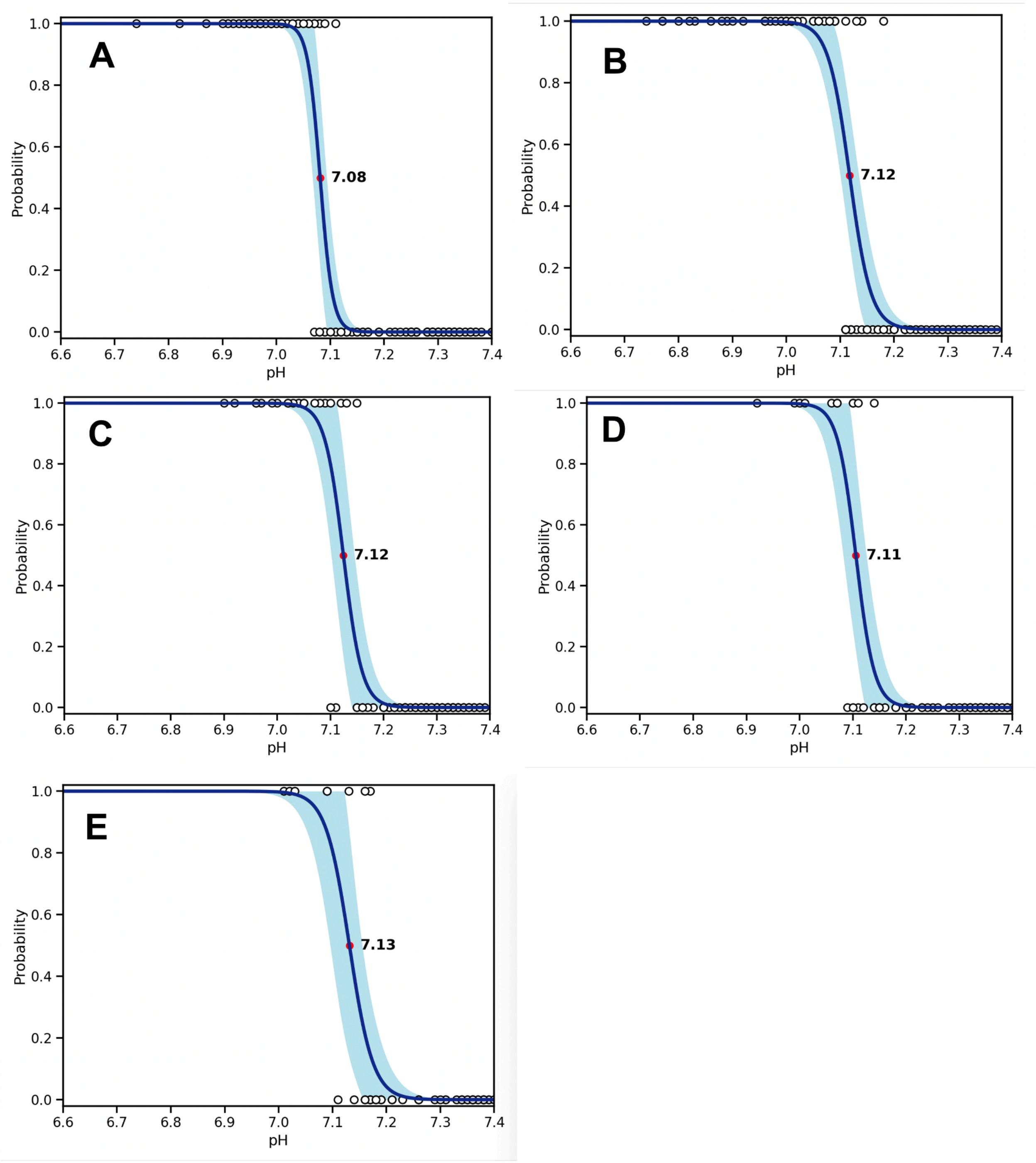


**Supplemental Figure S10.** Logistic regression fit of binary selected/unselected data with pH at discrete times points. (**A**) 0 hour (**B**) 1 hour (**C**) 2 hours (**D**) 3 hours (**E**) 4 hours.

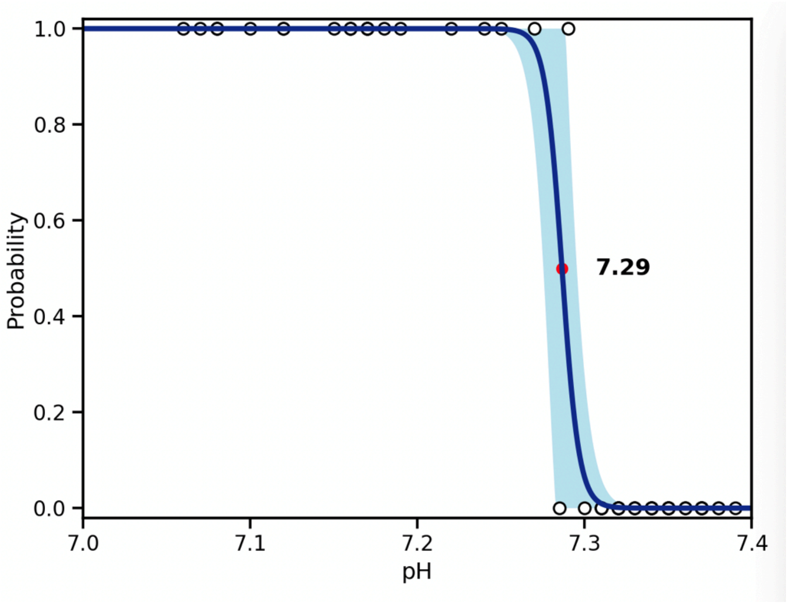


**Supplemental Figure S11.** Logistic regression fit of binary selected/unselected data with pH for activated Jurkat T cells.

**Supplemental Table S1. Typical flow parameters**. Channel geometry is provided below for reference. Negative flows below are opposite in direction to the main flow in the channel.

| Inlets and Outlets | Flow Rates (μL/min) |
|---|---|
| Aqueous Inlet | 0.3 |
| Oil Inlet | 3 |
| QX100 Inlet | 10 |
| Oil Entrainment Inlet | 45 – 49 |
| Oil Outlet | - 2 to -2.8 |

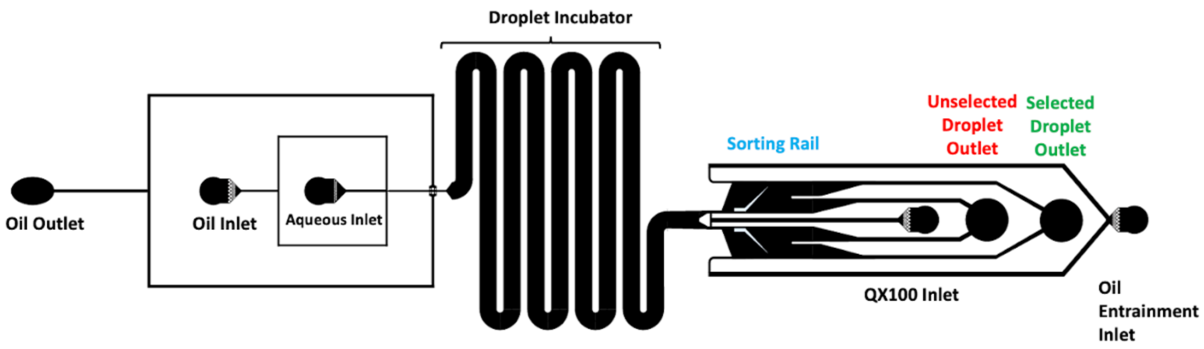
